# Soil microbial communities with greater investment in resource acquisition have lower growth yield

**DOI:** 10.1101/455071

**Authors:** Ashish A. Malik, Jeremy Puissant, Tim Goodall, Steven D. Allison, Robert I. Griffiths

**Affiliations:** Centre for Ecology and Hydrology, Wallingford, UK; Department of Ecology and Evolutionary Biology, University of California, Irvine, USA; Department of Earth System Science, University of California, Irvine, USA

## Abstract

Resource acquisition and growth yield are fundamental traits of microorganisms that have consequences for ecosystem functioning. However, there is a lack of empirical observations linking these traits. Using a landscape-scale survey of temperate near-neutral pH soils, we show tradeoffs in key community-level parameters linked to these traits. Increased investment into extracellular enzymes was associated with reduced growth yield; this reduction was linked more to carbon than nitrogen acquisition enzymes suggesting smaller stoichiometric constraints on community metabolism in examined soils.

---

Microorganisms are known to affect biogeochemical cycling of elements with consequences for ecosystem functioning. Of particular interest is how microbial metabolic strategies affect the fate of plant carbon entering soils^1,2^. Soil microorganisms partition the detrital carbon into biomass production and respiration, and this partitioning is key in determining the amount of carbon storage in soil^3,4^. Microbial growth yield, synonymously referred to as carbon use efficiency (CUE), determines the fraction of carbon that is allocated to biosynthetic processes (eg. growth) versus the fraction that is respired for cellular energy requirements^5,6^. Thus, growth yield integrates microbial physiology and is a measure of the energetic and material costs for survival and growth. Resource limitation can reduce growth yield by increasing the investment into metabolic machinery to degrade and take up complex substrates^7-9^. This investment to acquire energy- and nutrient-rich molecules comes in the form of extracellular enzymes and uptake proteins. Although there is some theoretical support to verify tradeoffs in growth versus resource acquisition, empirical validation of these tradeoffs in soil microbial communities is lacking. Nutrient limitation, particularly nitrogen, can also affect growth yield as cells need to maintain the elemental stoichiometry of their biomass^5,10,11^. Under such conditions, microbes may take up substrates in excess to meet nutrient requirements, leading to overflow respiration. Thus, it is also crucial to resolve the energetic and stoichiometric constraints on microbial growth yield in soil environments^12^.

We hypothesised that due to resource constraints, community-level tradeoffs exist between growth yield and resource acquisition, and that nutrient constraints affect community metabolism and reduce growth yield. To test this hypothesis, we assessed the empirical relationships between key physiological traits of soil microbial communities sampled at a landscape scale. Soil samples were collected in triplicate from 56 sites across Britain with land uses ranging from more pristine species-rich grasslands to intensive croplands that form a gradient of soil organic carbon concentration. However, here we focus on the physiology of communities in near neutral pH soils (38 sites), excluding those from acidic (pH < 6.2) wet soils that exhibited very slow growth rates and a peculiar metabolism (Supporting Information, figure S1-2). Resource acquisition traits were quantified by assessing the potential activities of the extracellular enzymes ß-1,4-glucosidase (BG), acetyl esterase (AE), leucine aminopeptidase (LAP) and ß-1,4-N-acetylglucosaminidase (NAG); BG and AE were used as a proxy for C acquisition and LAP and NAG were used as a proxy for N acquisition. Microbial growth yield was estimated as community CUE by tracing ^13^C-labelled, plant-derived substrates into total microbial DNA and respired CO_2_.

We observed a negative relationship between community CUE and potential investment in C acquisition (Figure 1a). BG catalyses the hydrolysis of glucose from cellobiose and AE is involved in deacetylation of xylans^12,13^. Both cellulose and hemicellulose, targets of the two enzymes, do not contain N and hence can be used as a proxy for C acquisition. Whereas, LAP catalyses the hydrolysis of proteins and NAG is involved in the hydrolysis of chitin and peptidoglycan, these target compounds are principle sources of N for microorganisms^12,14^. LAP and NAG have thus been widely used as a proxy for N acquisition. A similar negative relationship was observed between community CUE and N acquiring enzyme investment (Figure 1b), and although statistically significant this relationship was weaker relative to that between CUE and C enzyme investment. The distribution of these traits across the landscape was also related to the soil organic carbon (SOC) concentration gradient (overlaid in figure 1a-c). We have previously observed, in the same set of soils, that community CUE and biomass is positively related to SOC concentration (linear regression R^2^=0.34)^15^. Here we show that decreasing SOC was also linked to increasing C enzyme production rates (R^2^=0.3), and to a small extent increasing N enzyme investment (R^2^=0.06). These patterns suggest that in soils with lower SOC (usually intensive grasslands and arable croplands), resource limitation drives microbial communities to invest heavily into resource acquisition traits that trades off against growth yield. On the other hand, communities grow more efficiently in more resource-rich soils with higher SOM (usually “pristine” or less intensive grasslands) as they have lower investment in extracellular enzyme production and possess alternative substrate uptake mechanisms like ABC transporters^15,16^. It is also interesting to note that certain communities exhibited lower enzyme production and lower growth yield skewing the regression trends into non-linear functions (Figure 1a-b).

**Figure 1:**
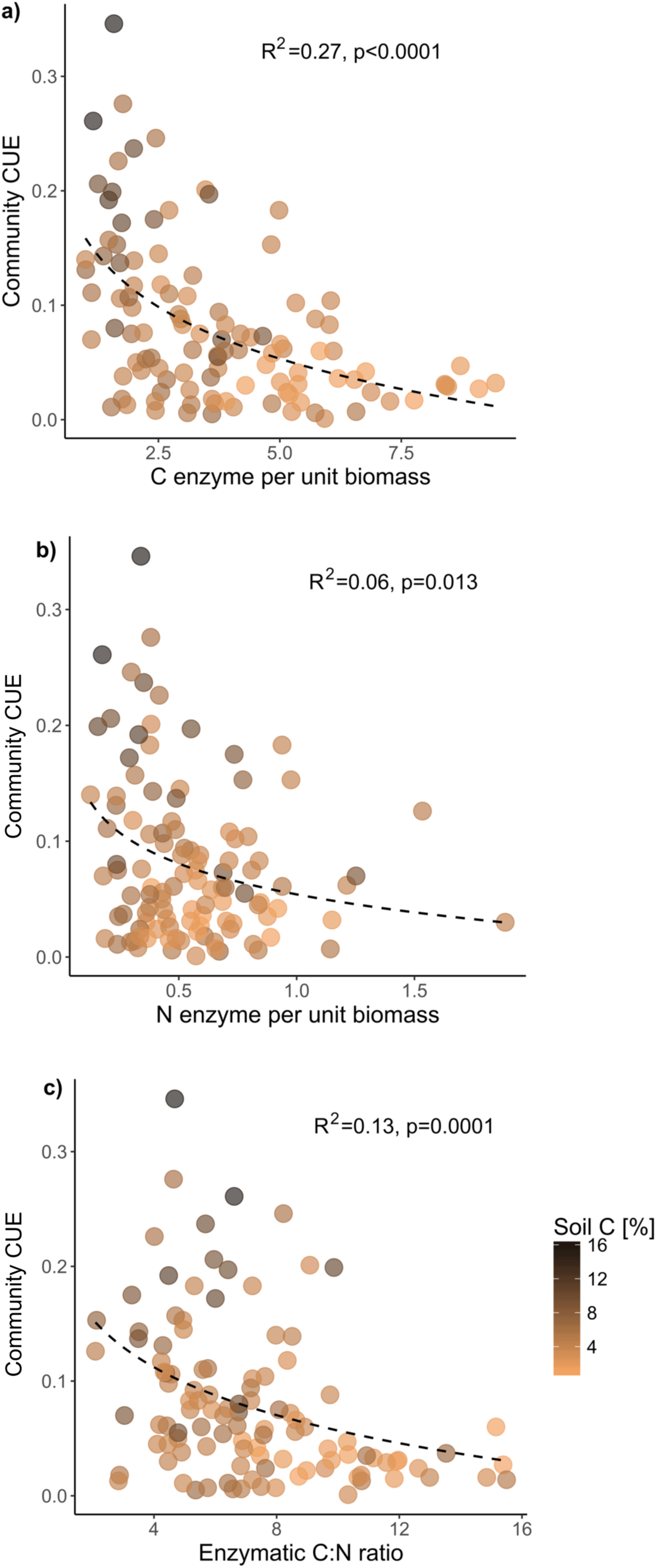
**(a-b)** Regression trends of community-aggregated growth yield or carbon use efficiency-CUE (unitless) with biomass specific C and N acquiring enzyme activity expressed as nmol min^−1^ μg-biomass-C^−1^ (DNA as a biomass proxy); **c)** Relationship between growth yield and ecoenzymatic C:N ratio. Overlaid on the scatterplots (a-c) is the variation in soil C concentration. Best fitting regression models were: a) y = −0.07ln(x) + 0.16; b) y = −0.03ln(x) + 0.06; c) y = −0.06ln(x) + 0.19.

Moreover, although the ecoenzymatic C:N ratio increases with decreasing CUE as we hypothesised (Figure 1c), there was little evidence to suggest stoichiometric constraints on microbial growth and metabolism. The stronger association of C- relative to N-acquiring enzyme production with CUE suggests that community-level energetic constraints are greater than stoichiometric constraints^12,17^. Still, this result could also reflect the resource and nutrient status of the temperate soils under investigation, which appeared to be C- and not N-limited. We also observed that ecoenzymatic C:N ratio and soil C:N ratio did not covary (Figure S3) indicating that soil C:N ratio is not a good indicator to link community physiology to their chemical environment^17^.

Based on empirical relationships, we provide evidence for a clear tradeoff between community-level growth yield and resource acquisition potential in near neutral pH soils. Although the statistical power of these relationships is not strong (given the geographically distributed nature of this survey), the patterns in trait distribution demonstrate distinct life history strategies. On the basis of these patterns, we applied a three-way microbial trait framework similar to Grime’s C-S-R triangle for plants^18^. Growth yield suffered in communities investing in maintenance requirements like resource acquisition through regeneration of extracellular enzymes (Figure 2, lower right). This tradeoff is reiterated by the absence of scenarios of communities excelling in both traits (figure 2, upper right). However, in certain communities lower resource acquisition costs were accompanied with lower growth yields (Figure 2, lower left), where, it is plausible that either or both of these traits trade off with some other unmeasured trait, likely stress tolerance^15,19,20^. In support of this interpretation, we previously found lower growth rate and yield in acidic soils (Figure S1-2)^15^ highlighting much higher maintenance costs of acid stress tolerance in such soils. Thus, we demonstrate strong support for the growth-maintenance tradeoff hypothesis and show trait tradeoffs have consequences for soil carbon dynamics.

**Figure 2:**
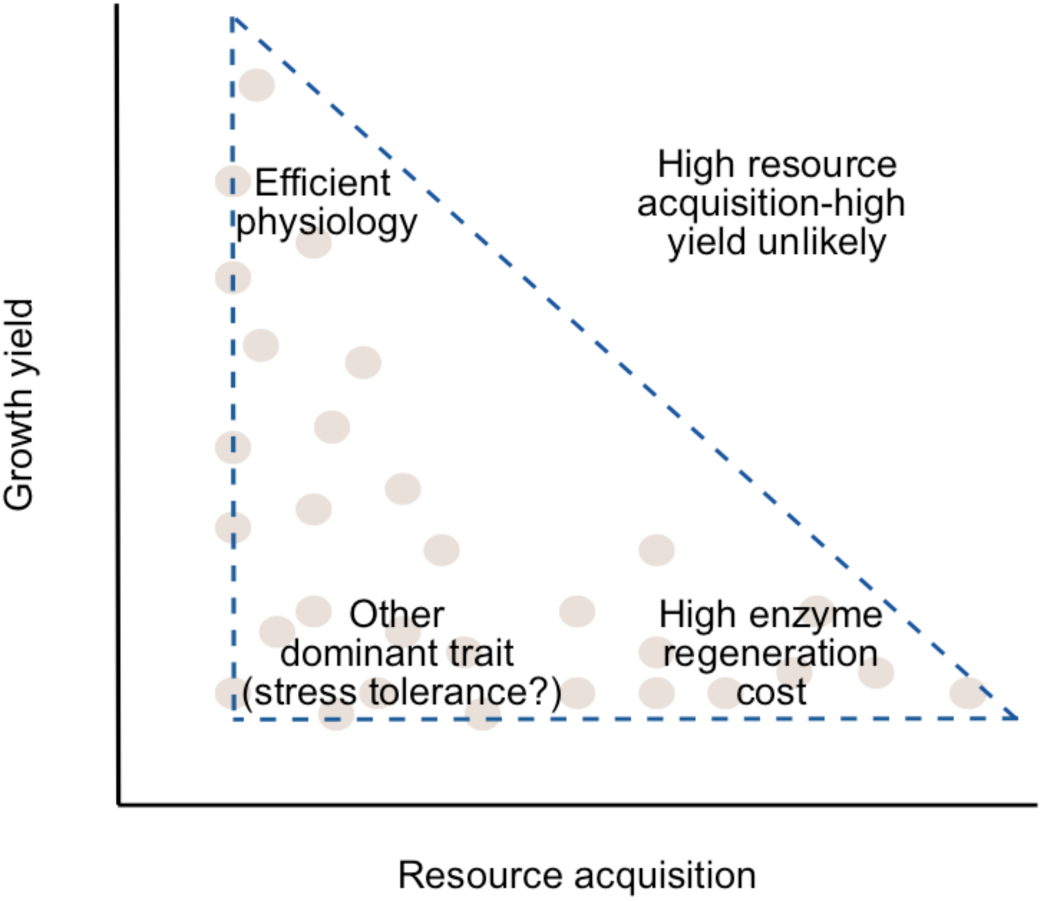
Conceptual framework assigning dominant life history strategies to microbial communities superimposed on the observed trait distribution patterns. Strategies for high yield, resource acquisition and stress tolerance are key traits that occupy the hypothesised microbial trait triangle.

In line with the empirical trends, we propose a microbial Y-A-S (high yield-resource acquisition-stress tolerance) life history 3 framework^21^, which suggests that tradeoffs in resource allocation among traits linked to high yield, resource acquisition and stress tolerance prevent microbes from excelling at multiple strategies such that different strategies are favoured under different environmental conditions. However, more work is required in estimating trait values for stress tolerance strategies and how they trade off with microbial growth yield. We also show, in the temperate soils under study, that stoichiometric imbalances have smaller impacts on microbial community growth yield in comparison to energetic requirements. This finding suggests that C flow in cellular systems is a fundamental constraint on microbial growth efficiency that affects the fate of plant and soil organic carbon.

## Methods

Here we used a subset of samples from a bigger soil survey, excluding low pH soils that exhibited very different microbial community physiology. From each of the 38 geographically distributed sites, 3 dispersed soil cores (5 cm diameter, 15 cm deep) were sampled. After all visible roots were removed, aliquots of the homogenized soil were used for the following functional analyses.

For microbial respiration measurements, a soil aliquot (1 g) was placed in a 10 mL glass vial, 100 µL of ^13^C-labeled plant leaf litter DOC solution (0.13 mgC) was added and incubated overnight (for ~16 h) in the dark at room temperature (21°C). Respired ^13^CO_2_ collected in the headspace was measured using a gas chromatography isotope ratio mass spectrometer (GC-IRMS, Delta+ XL, Thermo Fisher Scientific, Germany) coupled to a PAL-autosampler (CTC Analytics) with general purpose (GP) interface (Thermo Fisher Scientific, Germany).

Soil microbial total DNA was used as a proxy for biomass; DNA extraction was carried out on a soil aliquot of 0.25 g using PowerSoil-htp 96-well soil DNA isolation kit following manufacturer instructions (MO BIO Laboratories, UK). Another set of identical DNA extraction was performed following addition of 25 µL of the DO^13^C solution and overnight (16 h) incubation in dark. Both extracts were analysed in the size exclusion chromatography (SEC) mode on a liquid chromatography isotope ratio mass spectrometer LC-IRMS (HPLC system coupled to a Delta+ XP IRMS through an LC IsoLink interface; Thermo Fisher Scientific, Germany)^22^. This allowed us to obtain DNA-C content and the proportion of DO^13^C in microbial DNA. Microbial CUE was estimated as DNA-^13^C/(DNA-^13^C+∑CO_2_-^13^C), where ∑CO_2_-^13^C is the cumulative DO^13^C lost during respiration.

Activity of the extracellular enzymes was estimated with the common assay protocol using fluorigenic substrates^23^. ß-1,4-glucosidase, acetyl esterase, leucine aminopeptidase and N-acetyl glucosaminidase were assayed. Briefly, we homogenized 1.5 g soil in 20 ml of deionized water. The resultant slurry was used to perform enzyme activity assays using methylumbelliferyl (MUF) and 7-amino-4-methylcoumarin (AMC) conjugated substrates. The reaction was performed for 3 hours at 28°C, with one fluorometric measure every 30 minutes (BioSpa 8 Automated Incubator). Fluorescence intensity was measured using a Cytation 5 spectrophotometer linked to the automated incubator.

Visualisations and statistical analyses were performed with R software 2.14.0^24^. Regression models were run to assess empirical relationships between physiological traits of microbial communities.

## Acknowledgements

AAM has received funding from the European Union’s Horizon 2020 research and innovation programme under the Marie Skłodowska-Curie grant No 655240. We also acknowledge the UK Natural Environment Research Council for the Soil Security grant NE/M017125/1 to RIG, JP and TG, and the US DOE Genomic Science Program, BER, Office of Science project DE-SC0016410 funding to AAM and SDA.

## Author contributions

AAM, JP and RIG designed research; AAM and TG performed the stable isotope analyses; JP and TG performed the enzyme assays; AAM and JP performed statistical analyses; AAM and SDA developed the conceptual framework, AAM drafted the manuscript and all authors were involved in critical revision and approval of the final version.

